# Efficient Viral Capture and Inactivation from Bioaerosols Using Electrostatic Precipitation

**DOI:** 10.1101/2023.02.19.529105

**Authors:** Hannah E. Preston, Rebecca Bayliss, Nigel Temperton, Martin Mayora Neto, Jason Brewer, Alan L Parker

**Affiliations:** Division of Cancer and Genetics, Cardiff University School of Medicine, Heath Park, Cardiff, CF14 4XN, UK; Viral Pseudotype Unit, Medway School of Pharmacy, University of Kent, Central Avenue, Chatham, ME4 4BF, UK; Alesi Surgical Ltd, Medicentre, Heath Park Way, Cardiff, CF14 4UJ, UK; Systems Immunity University Research Institute, Cardiff University School of Medicine, Heath Park, Cardiff, CF14 4XN, UK

**Keywords:** Electrostatic Precipitation, Virus, Capture, Inactivation, Adenovirus, SARS-CoV-2

## Abstract

The presence of infectious viral particles in bioaerosols generated during laparoscopic surgery places surgical staff at significant risk of infection and represents a major cause of nosocomial infection. These factors contributed to the postponement and cancellation of countless surgical procedures during the early stages of the ongoing COVID-19 pandemic, causing backlogs, increased waiting times for surgical procedures and excess deaths indirectly related to the pandemic. The development and implementation of devices that effectively inactivate viral particles from bioaerosols would be beneficial in limiting or preventing the spread of infections from such bioaerosols. Here, we sought to evaluate whether electrostatic precipitation (EP) is a viable means to capture and inactivate both non-enveloped (Adenovirus) and enveloped (SARS-CoV-2 Pseudotyped Lentivirus) viral particles present in bioaerosols. We developed a closed-system model to mimic the release of bioaerosols during laparoscopic surgery. Known concentrations of each virus were aerosolised into the model system, exposed to EP using a commercially available system (Ultravision^TM^, Alesi Surgical Limited, UK) and collected in a BioSampler for analysis. Using qPCR to quantify viral genomes and transduction assays to quantify biological activity, we show that both enveloped and non-enveloped viral particles were efficiently captured and inactivated by EP. Both capture and inactivation could be further enhanced when increasing the voltage to 10kV, or when using two Ultravision^™^ discharge electrodes together at 8kV. This study highlights EP as an efficient means for capturing and inactivating viral particles present in bioaerosols. The use of EP may limit the spread of diseases, reducing nosocomial infections and potentially enable the continuation of surgical procedures during periods of viral pandemics.

**Highlights:**

- Bioaerosols released from patients during surgery have the potential to facilitate viral spread.
- Ultravision^™^ technology works via the process of electrostatic precipitation.
- Electrostatic precipitation can be manipulated to capture and inactivate aerosolised viral particles, preventing viral spread.
- Electrostatic precipitation is effective against both enveloped and non-enveloped viral particles.
- Electrostatic precipitation represents a viable means to reduce nosocomial infections.

**Graphical Abstract:** 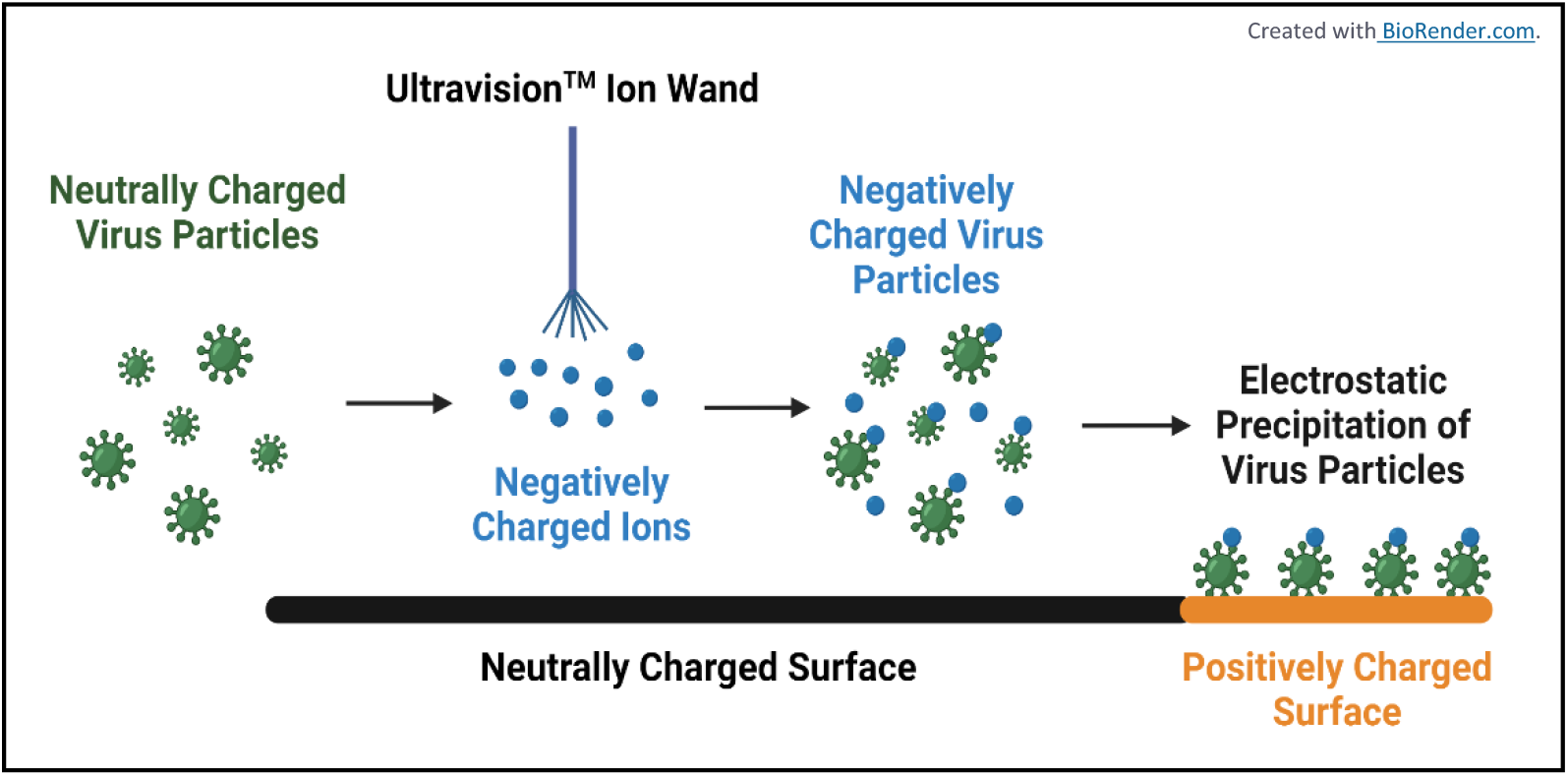

## 1. Introduction

Acute respiratory viruses are the fourth leading cause of mortality worldwide [1]. Although respiratory viruses can be spread by physical contact, contaminated fomites, and large droplets, key transmission occurs via the dispersion of bioaerosols from an infectious individual [2]. Additionally, previous studies have shown that wildtype non-respiratory viruses, such as Human Immunodeficiency Virus (HIV) and Human Papillomavirus (HPV) can also be released in bioaerosols, during aerosol-generating medical procedures, enabling viral transmission [3, 4].

With particular focus on the 2019 SARS-CoV-2 pandemic, >640 million cases and >6.5 million directly related deaths were reported worldwide in December 2022 [5]. Regarding the indirect consequences of the pandemic, it is estimated that hundreds of thousands of surgeries were delayed or cancelled as a result. Bioaerosol-generating procedures, including laparoscopy, tracheostomy, open suctioning, and administration of nebulised treatments were at the highest risk of cancellation, due to the likelihood of airborne transmission to staff and other patients [6]. This has left patients untreated and undiagnosed, creating enormous backlogs of waitlisted surgeries, thereby increasing the demand for private health care [7].

Mitigation strategies such as mask wearing, personal protective equipment (PPE), social distancing, isolation of infected patients and mass vaccinations were enforced and encouraged by the health authorities to reduce the spread of SARS-CoV-2 [8]. However, cases of SARS-CoV-2 infection continue to fluctuate at high levels, due to the evolution of new viral strains, easing of government-enforced restrictions and a lack in vaccine confidence by the general public [9, 10]. Therefore, the population remains at risk, emphasising the need for novel non-pharmaceutical interventions (NPIs).

Commonly used NPIs for reducing the spread of disease in hospitals are Ultra-Low or High-Efficiency Particulate Absorbing (ULPA, HEPA) filters and Ultraviolet (UV) light sterilisation [11]. Although these NPIs are somewhat capable of purifying indoor air, each system is hindered by limitations. ULPA/HEPA filters are non-economical and labour intensive, as they use high levels of energy to run and require regular filter changes. Further, most filters trap particles 12μm in diameter, yet many viruses, including SARS-CoV-2, are smaller than this [12]. Viruses that are trapped via a filter can remain live and active, adding an additional risk to their use within hospitals and requiring appropriate treatment as a biohazard during disposal [12]. UV light is capable of inactivating viruses, however its efficiency is limited to its alignment with and distance from the virus itself [13]. As well as this, the exposure time and irradiance doses of UV light used to decontaminate indoor environments has not been well standardised, and incorrect usage of UV light can be hazardous [13].

As nosocomial virus transmission occurs most commonly by the release of bioaerosols from infectious patients, it would be beneficial to develop a NPI that efficiently captures and inactivates viral particles from bioaerosols in hospital environments. Ultravision^™^ technology has been developed to be used during key-hole surgeries, such as abdominal laparoscopies, to eliminate surgical smoke [14]. Surgical smoke is produced by the thermal destruction of tissue by electrosurgical instruments during medical procedures and can obstruct the surgeons field of vision, resulting in safety implications [15]. Surgical smoke consists of 95% water vapor and 5% cellular debris, of which can contain live bacterial and viral particles [15]. Ultravision^™^ clears surgical smoke by using electrostatic precipitation (EP), which involves the generation of an electric field to precipitate particles out of aerosolised suspension and onto a collection surface [16]. The Ultravision^™^ Ionwand functions as a discharge electrode, by emitting negatively charged ions into a neutrally charged space, thus creating a corona discharge [17]. The negative current produced from the Ultravision^™^ Ionwand results in the creation of low-energy gas ions and subsequent transient electrostatic charging of aerosolised matter within a local atmosphere. In addition, a return electrode carrying a positive charge is connected to a collector plate, enabling the precipitation of negatively charged particles via electrostatic attraction. This mechanism is exploited during key-hole surgery to clear surgical smoke, whereby aerosolised particles are ionised by the Ultravision^™^ Ionwand and precipitated onto the patient’s abdominal tissue, which is connected to a positively charged return electrode pad [18].

It has been suggested that EP could be used in point-of-care systems as a method of aerosol sampling, to diagnose patients rapidly and accurately for respiratory viral infections, reducing the need to perform invasive and uncomfortable diagnostic procedures such as bronchoscopy [19]. Furthermore, EP has been incorporated into a microfluidic lab-on-chip device, for immediate pathogenic detection from aerosol droplets released in the exhaled breath of patients [19]. Custom bioaerosol samplers, employing EP mechanisms have also been developed and demonstrated to detect airborne Influenza Virus particles; of which studies have claimed may reduce sampling times down from hours to minutes, thus inhibiting viral transmission faster than currently existing approaches [20].

Since Ultravision^™^ functions to clear surgical smoke via EP, it was rational to evaluate the ability of EP to capture and inactivate aerosolised viral particles using Ultravision^™^. Furthermore, Ultravision^™^ has already been cleared by regulators as safe and effective in use [21, 22], thereby serving as a practical, multi-modal device to use during medical procedures to prevent the spread of aerosolised viral particles. In addition, EP is capable of precipitating particles at a minimum diameter of 7nm [23], thus improving the efficiency of particle capture and filtration compared to other established and commonly used ventilation and filtration systems.

The objective of our study was to evaluate the capture and inactivation of bioaerosol-containing viral particles by EP. Non-enveloped (Ad5) and enveloped (SARS-CoV-2 Pseudotyped Lentivirus) viral particles were aerosolised into a closed-system model, that was representative of key-hole surgery, and exposed to EP. Recovered samples were analysed for viral presence by qPCR of viral genomes and for biological activity by transduction and plaque assays in target cell lines. We hypothesised that viral exposure to EP would result in significant viral capture and inactivation.

Reducing viral transmission is not limited to SARS-CoV-2, but accounts for all viral outbreaks that may lead to future pandemics. It is therefore important that novel NPI’s are evaluated and developed, to increase our preparation, improve safety within hospitals and prevent the need to cancel surgeries and medical procedures in the case of future pandemics.

## 2. Methods & Materials

### 2.1 Production of Viruses

Ad5 was modified to express GFP and was propagated in T-REx-293 cells expressing E1 gene products and purified using Caesium Chloride gradient ultracentrifugation as previously described [24]. Stock titres were determined by Micro-BCA assay (Pierce, Thermo Fisher, Loughborough, UK), assuming that 1μg protein was equal to 4 x 10^9^ virus particles (vp) and monodispersity was confirmed by Nanoparticle Tracking Analysis (NanoSight NS300, Malvern, UK). Infectious titres were quantified by end-point dilution plaque assay, performed in T-REx-293 cells, determining plaque forming units per millilitre (PFU/ml).

The SARS-CoV-2 Pseudotyped Lentivirus (SARS-2 PV) contained a HIV core and expressed Wuhan strain SARS-CoV-2 Spike Proteins (GenBank accession: 43740568) on their viral envelope. SARS-2 PV are replication deficient and express GFP under the control of a SFFV promoter post transduction [25, 26]. SARS-2 PV were produced in HEK-293T/17 cells (ATCC CRL11268) that were pre-seeded in a T175 flask (Thermo) with approximately 5 x10^6^ cells the day before transfection. Cells were then co-transfected with 2 μg of packaging lentiviral core p8.91, 3 μg of pCSGW encoding Green Fluorescent Protein, and 2 μg of the spike SARS2 (D614G)-pCAGGS (Medicines & Healthcare Products Regulatory Agency CFAR100985) using FugeneHD (Promega) transfection reagent at a ratio of 1:3 DNA:Fugene in optiMEM (Gibco). SARS-2 PV were harvested at 48h post transfection and supernatant filtered through a 0.45 μm acetate cellulose filter (Starlab) [27] [28]. Functional titres were determined by plaque assay.

### 2.2 Cell Lines

T-REx-293 (Tetracycline Repressor Protein expression: Invitrogen^TM^ R71007) and HEK-293T cells (Human Embryonic Kidney: ATCC CRL-1573) were used to produce Ad5 and SARS-2 PV virus stocks, respectively. CHO-CAR (Chinese Hamster Ovarian cells, transfected to express Human CAR) and CHO-ACE2-TMPRSS2 stable line (Chinese Hamster Ovarian cells expressing Human ACE2 and TMPRSS2) cells were used in transduction assays with Ad5.GFP and SARS-2 PV, respectively. T-REx-293 and HEK-293T cells were cultured in DMEM media (Dulbecco’s Modified Eagle’s Medium; Sigma-Aldrich, Gillingham, UK #D5796), whilst CHO-CAR and CHO-ACE2-TMPRSS2 cells were cultured in DMEM-F12 media (Dulbecco’s Modified Eagle’s Medium/Nutrient Mixture F-12 Ham; Sigma-Aldrich, Gillingham, UK #D0697). All media was supplemented with 10% FBS (Foetal Bovine Serum; Gibco, Paisley, UK #10500-064), 2% Penicillin and Streptomycin (Gibco, Paisley, UK #15070-063) and 1% L-Glutamine (stock 200 mM; Gibco, Paisley, UK #25030-024). CHO-ACE2-TMPRSS2 cells were also passaged with 2μg/mL Puromycin and 100μg/mL Hygromycin once a week. Cells were grown at 37°C with 5% CO_2_. Dulbecco’s Phosphate Buffered Saline (PBS, Gibco^TM^, #10010023) and 0.05% Trypsin (Gibco^TM^, #11590626) were used for subculture.

### 2.3 Experimental Setup of the Closed-System Model

The standard closed-system model (***Figure 1***) was optimised and altered for some experiments, however the general setup remained consistent in each run. A medical grade nebuliser (Aerogen^®^ Solo Starter Kit, Aerogen Ltd, Galway, AG-A53000-XX) was used to aerosolise 10ml of each sample into a 3L reaction kettle (QuickFit^TM^ Wide Neck Flask Reaction 3L, Scientific Laboratory Supplies Ltd, UK, QFR3LF). The reaction kettle was fitted with a lid containing multiple culture vessels (QuickFit^™^ Borosilicate Glass Flange Lid, Fisher Scientific, Leicestershire, MAF3/52), enabling the insertion of samples and materials, whilst maintaining an air-tight system. The Ultravision^TM^ power supply (Ultravision^TM^ Generator, BOWA Medial UK, Newton Abbot, DAD-001-015) was stationed outside of the closed system. The Ultravision^™^ Ionwand discharge electrode (Ionwand^™^, BOWA Medial UK, Newton Abbot, DAD-001-003) was inserted into the reaction kettle through a Suba-Seal^®^, 15cm from the bottom of the reaction kettle and 7cm from either side of the reaction kettle. The Ultravision^™^ generator was attached to copper tape that covered the inside of the reaction kettle, using a modified patient return electrode cable, functioning as a positively charged collector-plate. Stopcock adapters (QuickFit^™^ Borosilicate Glass Stopcock Adaptors with Sockets, Fisher Scientific, Leicestershire, MF14/3/SC) were placed throughout the system, ensuring unidirectional flow of the aerosol. A vacuum unit (Duet Flat-Back Aspirator, SSCOR, US, 2314B) was used, at maximum flow rate, to suction the aerosol through the system and into a BioSampler (BioSampler^®^, SKC Ltd, Dorset, 225-9595). The BioSampler (assembled as per manufacturer’s instructions) contained 2ml sterile serum-free media (DMEM) to recover the captured aerosol samples. To prevent viral contamination, a cold-trap (QuickFit^™^ Cold-trap, VWR, Pennsylvania, 201-3052) was fitted between the BioSampler and the vacuum unit. All experimentation was conducted in a Class II laminar flow hood, and all materials were autoclaved or sterilised with 70% Industrialised Methylated Spirit (IMS) before and after use.

**Figure 1.**
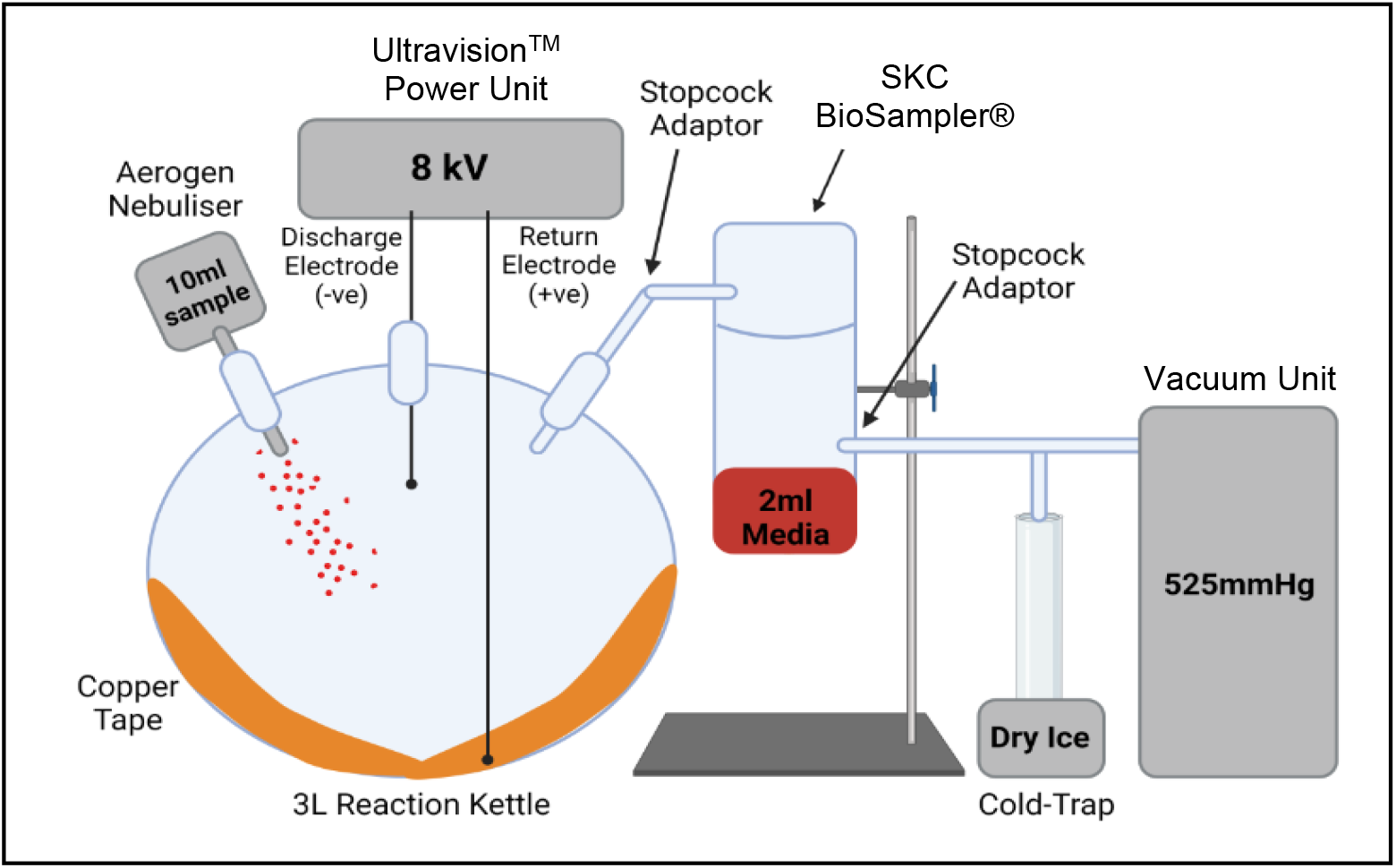
Schematic Depicting the Experimental Setup of the Refined Closed-System Model. All samples were aerosolised into the air-tight reaction kettle, exposed to Ultravision^™^ (active/inactive) and suctioned into the BioSampler for recovery and collection. Collected samples were stored at −80°C immediately after each experimental run, prior to experimental analysis.

### 2.4 Experimental Protocol

In each experimental run, 10ml samples were aerosolised into the reaction kettle, which was heated to 37°C, and exposed to an inactivate/active Ultravision^TM^ Ionwand (turned off/turned on), until the entire sample had been completely aerosolised. The samples were suctioned into the BioSampler, where they were recovered in 2ml serum-free media (DMEM). Samples collected in the BioSampler were stored at −80°C immediately after each run, in preparation for experimental analysis. Serum-free media was aerosolised through the system as a negative control, in each run. Additionally, 2ml samples of Ad5.GFP at 1×10^10^vp/ml and neat SARS-2 PV (at approximately 3 x 10^7^pfu/ml) that were not aerosolised, nor exposed to the system, were stored at −80°C, to use as positive (untreated) controls. For each experimental run, Ad5.GFP was diluted to 1×10^10^vp/ml, whilst SARS-2 PV samples were aerosolised neat.

### 2.5 Quantification of Viral Genome Copy Number by qPCR

DNA was extracted using the QIAamp MinElute Virus Kit (Qiagen, #57704). Purified DNA was eluted in 50μl of Ultra-Pure Water (UltraPure^™^ DNase/RNase-Free Distilled Water, Invitrogen^™^, Thermo Fisher, #11538646) and stored at −20°C. DNA extracted from the virus stocks were used as standards (Serial dilution: undiluted (200ng/μl), 10^-1^, 10^-2^, 10^-3^, 10^-4^, 10^-5^ and 10^-6^). DNA extracted from experimental samples remained undiluted. Primers (Ad5 Hexon Forward: CCTGCTTACCCCCAACGAGTTTGA, Ad5 Hexon Reverse: GGAGTACATGCGGTCCTTGTAGCTC; P24 Capsid: Forward: GGCTTTCAGCCCAGAAGTGATACC, P24 Capsid Reverse: GGGTCCTCCTACTCCCTGACATG) were used at 10Mm. qPCR for viral DNA was performed using the SYBR Green Master Mix (PowerUp^™^ SYBR^™^ Green Master Mix, Applied Biosystems^™^, Thermo Fisher, #A25741) (per reaction: 15μl Master Mix and 5μl DNA). Reactions were performed in triplicate (for both samples and standards). QuantStudio^™^ software was used to set the thermal cycling conditions of the qPCR (Pharmaceutical Analytics QuantStudio^™^ 5 Real-Time PCR System, AppliedBiosystems^™^, Thermo Fisher, #A31670). Samples were held at 50°C for 2 min, followed by 95°C for 2 min. Samples were then cycled at 95°C for 15 sec and 60°C for 1 min for 40 cycles.

### 2.6 Transduction Assays

CHO-CAR/CHO-ACE2-TMPRSS2 cells were seeded into a 96-well plate at a density of 2×10^4^ cells/well in 200μl complete media and cultured overnight. The following day, complete media was removed, cells were washed briefly in PBS, and experimental samples were added to the cells (100μl, undiluted) and incubated at 37°C for 3 hours. The media was then removed and discarded, and the cells were washed twice with 100μl PBS, prior to replenishing the cells with 200μl total media and culturing for an additional 48 hours. Cells were visualised for GFP expression using an EVOS (EVOS M7000, Invitrogen^™^, Thermo Fisher Scientific, #AMF7000), then harvested in FACS buffer and fixed with 4% Paraformaldehyde. Flow Cytometry was performed, using the Accuri (BD Accuri C6 v.1.0.264.21) and the FL1-A channel, to detect virally transduced cells. FlowJo^™^v10 software was used to analyse all Flow Cytometry data.

### 2.7 Plaque Assays

T-REx-293/HEK-293T cells were seeded in 12-well plates in complete media, at a density of 1×10^5^ cells/well in triplicate. Cells were cultured for 24 hours, prior to the experiment. Media was removed, and the cells were washed with 1ml PBS. Experimental samples were added to the wells (1ml, undiluted) in duplicate. The cells were incubated at 37°C for 2 hours, then the media was removed and replaced with 1ml complete media. The cells were cultured for a further 48 hours, before analysis. EVOS microscopy (EVOS M7000, Invitrogen^™^, Thermo Fisher Scientific, #AMF7000) was used to image the cells (Objective Lens X20). Transduced cells fluoresced green light under the GFP light source, enabling manual counting of infected cells. The PFU/ml of each sample was calculated using the formula:

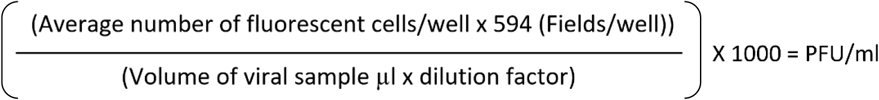

### 2.8 Statistical Analysis

All data presented show the mean ± SD. GraphPad Prism v4.03 (GraphPad Software Inc., La Jolla, CA) was used to produce all bar chart figures. The GraphPad Quickcalcs t-test calculator was used to perform the two-tailed paired t-test. p-Values of * = p<0.05, ** = p<0.005, *** = p<0.0005, ns = not statistically significant, p>0.05.

## 3. Results

To mimic the release of bioaerosols that occurs during key-hole surgery, we developed a closed-system model representing the peritoneal cavity and the Ultravision^™^ set up in a laparoscopic procedure. A 3L reaction kettle was used to resemble the peritoneal cavity, which is sufflated to approximately 3L with CO2 during laparoscopy [29]. The Ultravision^™^ Ionwand was positioned within the reaction kettle, directly above the region of bioaerosol release, as it would be during laparoscopy. Quick-fit^®^ glassware was used to ensure that the entire model was air-tight, preventing the release of virally contaminated aerosols. A vacuum unit was employed to suction the aerosol through the closed-system model in a unidirectional flow into a SKC BioSampler^®^ for sample recovery to assess viral presence within the aerosol, following exposure to Ultravision^™^. Recovered samples collected in the BioSampler were analysed for viral presence by qPCR and for viral activity via transduction and plaque assays. In addition, physical parameters thought to affect the efficiency of EP were altered, in an attempt to determine optimal settings for the usage of Ultravision^™^. Such parameters included temperature, voltage, the number of Ultravision^™^ Ionwands within the reaction kettle and the material of the collector plate attached to the positively charged return electrode. Further, two different virus samples were aerosolised and exposed to Ultravision^™^ in the closed-system model, to evaluate the ability of EP to capture and inactivate both non-enveloped (Ad5) and enveloped (SARS-PV) viral particles. Both viruses were rendered replication deficient and were genetically modified to express GFP, enabling detection in experimental assays.

### 3.1 Ad5 Particles were Successfully Captured and Inactivated by Electrostatic Precipitation when Aerosolised at 37°C

From preliminary experiments it was found that the closed-system model needed to be heated at 37°C to eliminate sample condensation. First, we sought to evaluate whether EP could capture and inactivate aerosolised non-enveloped Ad5 particles using our standard closed-system model. The number of recovered Ad5 genomes significantly decreased following Ad5 exposure to inactive EP as gauged by qPCR for viral genomes, indicating viral loss as a result of sample aerosolization alone ***(Figure 2.A***). A significant 6.8-fold reduction in the number of recovered Ad5 genomes was observed following Ad5 exposure to active EP ***(Figure 2.A***). Ad5 viability was not affected following exposure to inactive EP, as displayed by transduction and plaque assays ***(Figure 2.B & C***), indicating that sample aerosolization at 37°C was not detrimental to Ad5. The transduction assay demonstrated a 13.6-fold reduction in the percentage of transduction, in cells that were treated with Ad5 that had been exposed to active EP ***(Figure 2.B***). Mirroring this, the plaque assay displayed a 4×10^3^-fold reduction in active Ad5 particles, in the sample exposed to active EP ***(Figure 2.C & D***). These results indicated that EP successfully captured and inactivated aerosolised Ad5 particles within our standard closed-system model.

**Figure 2.**
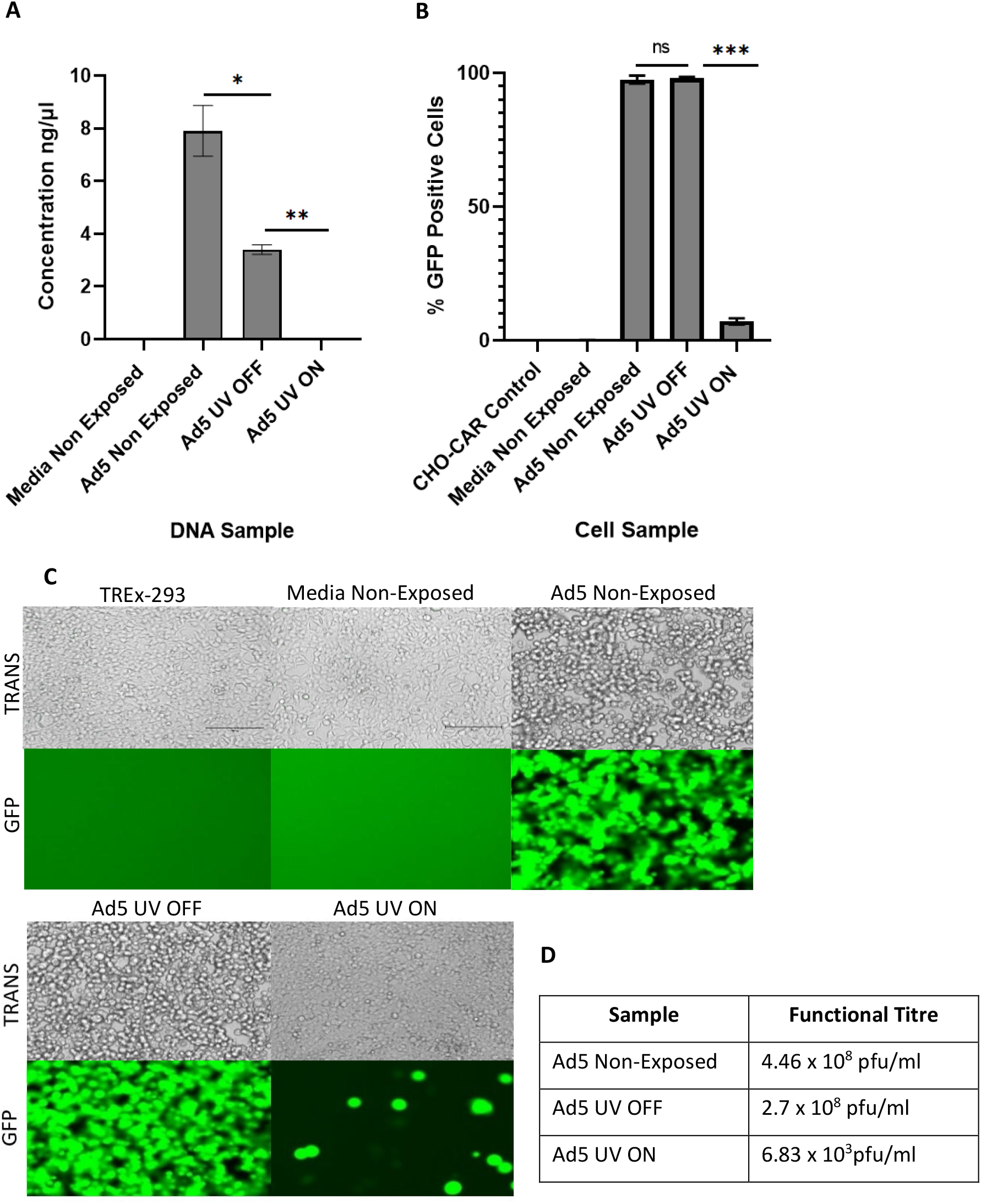
Capture and Inactivation of Ad5 by Electrostatic Precipitation. ‘UV OFF’ signifies sample exposure to inactive Ultravision^™^ and ‘UV ON’ signifies sample exposure to active Ultravision^™^. ‘Non-Exposed’ signifies samples that were not aerosolised through the model system, nor exposed to Ultravision^™^. **(A)** Viral capture quantified by qPCR. **(B)** Viral inactivation demonstrated by transduction assay. **(C & D)** Viral inactivation displayed by plaque assay in TREx-293 cells. TREx-293 cells treated with samples and analysed for GFP fluorescence. TRANS = Brightfield transmitted light, GFP = GFP light source. Error bars represent the ±SD (n = 3). Plaque assay functional titres represent the mean (n = 5).

### 3.2 Capture and Inactivation of Ad5.GFP was Most Efficient when Exposing Viral Particles to 10kV

Multiple parameters may impact the efficiency of EP. We assessed the impact of increasing voltages on the ability of EP to capture and inactivate aerosolised Ad5. Ultravision^™^ is currently used at 8kV to clear surgical smoke during laparoscopies. We exposed aerosolised samples of Ad5 to Ultravision^™^ active at 6kV, 8kV and 10kV, to determine whether decreasing or increasing the current voltage impacted its ability to capture and inactivate viral particles. As 10kV is the maximum voltage that is medically approved for Ultravision^™^ use during surgery, voltages above this were not evaluated.

qPCR analysis of treated samples indicated significant viral capture by EP, following sample exposure to 6kV, 8kV and 10kV (***Figure 3.A***). The number of viral genomes were reduced by 21.8-fold and 16.8-fold, following Ad5 exposure to 6kV and 8kV, respectively. However, Ad5 capture was enhanced when exposing the viral particles to 10kV, as shown by a 7.4×10^3^-fold reduction in the number of viral genomes ***(Figure 3.A***). Increasing the voltage to 10kV also improved viral inactivation, demonstrated by transduction and plaque assay ***(Figure 3.B & C***). The percentage of transduced cells infected with Ad5 samples that had been exposed to 6kV and 8kV was significantly reduced by 6.6-fold and 25.6-fold, respectively ***(Figure 3.B***). Cells treated with Ad5 that had been exposed to 10kV displayed a 529.4-fold reduction in viral transduction ***(Figure 3.B***). Mirroring this, plaque assays of treated samples demonstrated a significant decrease in the number of viable Ad5 particles in samples that were exposed to 6kV, 8kV and 10kV ***(Figure 3.C & D***). Imagining of GFP highlighted a complete absence of viable Ad5 particles in cells infected with Ad5 samples that had been exposed to 10kV, indicating that 10kV is the optimal voltage to elicit efficient EP of bioaerosols during surgery, to completely prevent the transmission of infectious aerosolised virus particles ***(Figure 3.C***).

**Figure 3.**
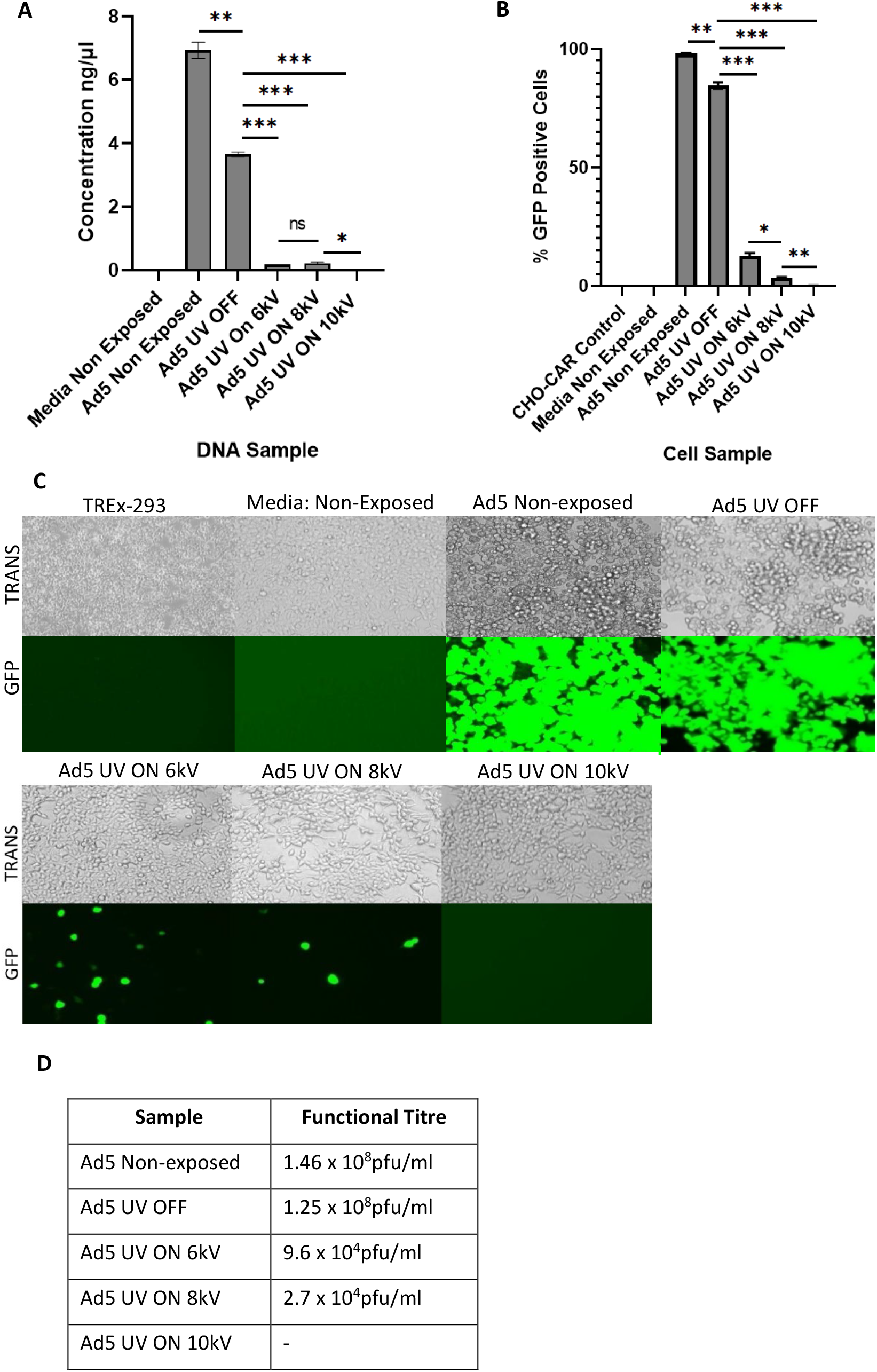
Increasing the Voltage of Ultravision^™^ to 10kV Enhances Viral Capture and Inactivation. ‘UV OFF’ signifies sample exposure to inactive Ultravision^™^ and ‘UV ON’ signifies sample exposure to active Ultravision^™^. ‘Non-Exposed’ signifies samples that were not aerosolised through the model system, nor exposed to Ultravision^™^. **(A)** Viral capture demonstrated by qPCR. **(B)** Viral inactivation determined by transduction assay. **(C & D)** Viral inactivation displayed by plaque assay in TREx-293 cells. TREx-293 cells treated with samples and analysed for GFP fluorescence. TRANS = Brightfield transmitted light, GFP = GFP light source. Error bars represent the ±SD (n = 3). Plaque assay functional titres represent the mean (n = 5).

### 3.3 Using 2 Ultravision^™^ Ionwands Enhanced Adenoviral Capture and Inactivation

We next evaluated whether enhanced viral inactivation was possible when exposing aerosolised Ad5 particles to 2, rather than a single, Ultravision^TM^ Ionwands. Both Ionwands were used at 8kV, maintaining the voltage setting that is currently used during laparoscopic surgery. Separate Ad5 samples were exposed to either 1 or 2 Ionwands, to evaluate whether combining 2 Ionwands improved viral capture and inactivation.

qPCR results displayed a significant decrease in the number of viral genomes in Ad5 samples that were exposed to either 1 or 2 active Ionwands. A 125-fold reduction in the number of Ad5 genomes was observed in the sample exposed to 1 active Ionwand, whereas exposure of Ad5 to 2 Ionwands resulted in an increased 1.25×10^3^-fold reduction in the number of Ad5 genomes detected ***(Figure 4.A***). This indicated that using 2 Ionwands, both active at 8kV, enhanced viral capture by a further 10-fold. Similarly, Ad5 samples exposed to 1 or 2 Ionwands were both significantly inactivated. Cells treated with the Ad5 sample that had been exposed to a single active Ionwand displayed a 31.6-fold reduction in the percentage of virally transduced cells ***(Figure 4.B***). In comparison, cells treated with the Ad5 sample that had been exposed to 2 active Ionwands displayed a 215.2-fold reduction in the percentage of transduced cells, indicating that using 2 Ionwands enhanced viral capture ***(Figure 4.B***). Plaque assay confirmed these findings, as shown by an 800-fold decrease in the number of active Ad5 particles, post exposure to a single Ionwand, in comparison to a complete elimination of active Ad5 particles, post exposure to 2 Ionwands ***(Figure 4.C & D***). This experimental run highlighted that using 2 Ionwands enhanced viral capture and inactivation in a synergistic manner.

**Figure 4.**
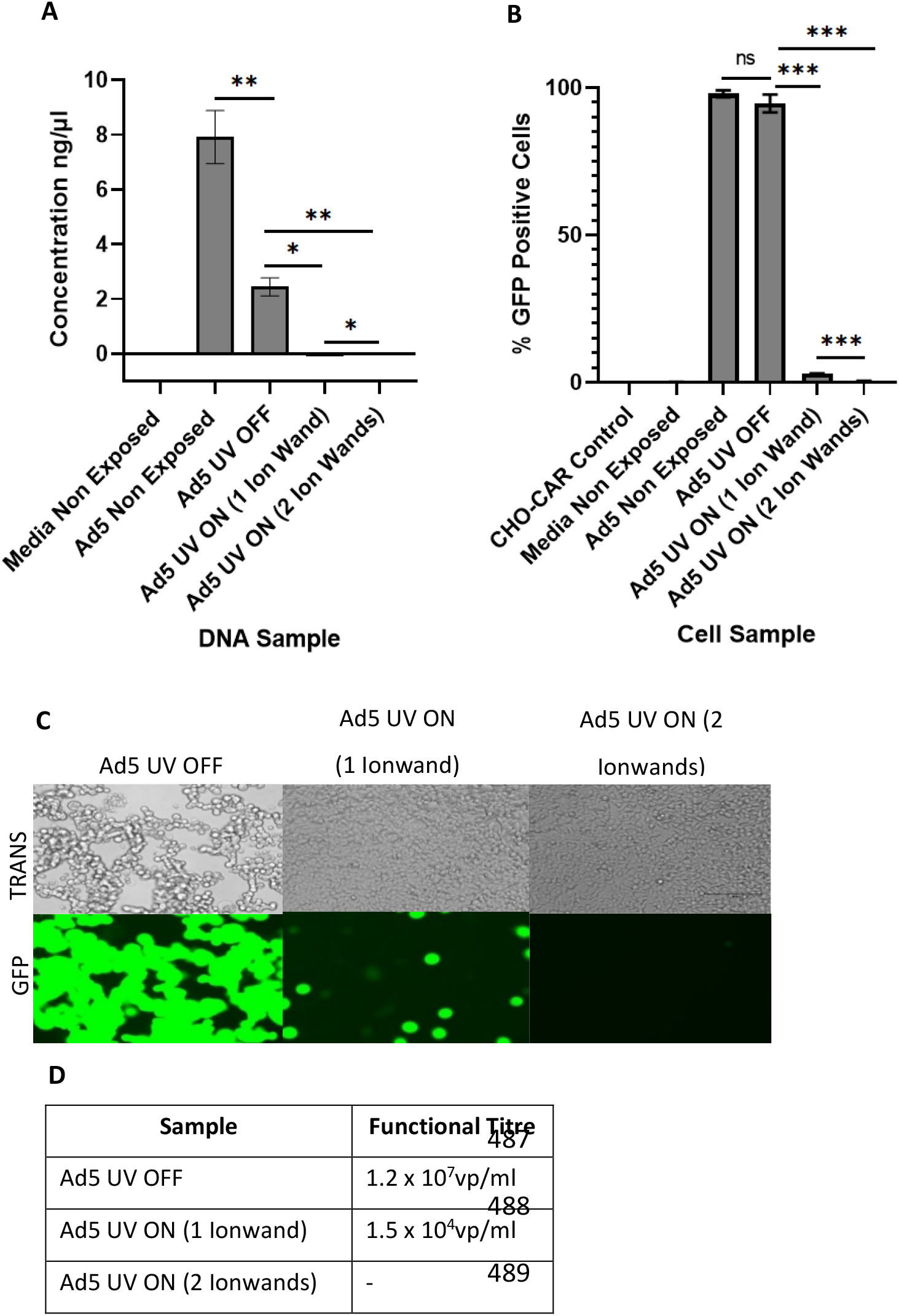
Exposing Ad5 Particles to 2 Ionwands, opposed to 1, Enhances Viral Capture and Inactivation. ‘UV OFF’ signifies sample exposure to inactive Ultravision^™^ and ‘UVON’ signifies sample exposure to active Ultravision^™^. ‘Non-Exposed’ signifies samples that were not aerosolised through the model system, nor exposed to Ultravision^™^. **(A)** Viral capture demonstrated by qPCR. **(B)** Viral inactivation determined by transduction assay. **(C & D)** Viral inactivation displayed by plaque assay in TREx-293 cells. TREx-293 cells treated with samples and analysed for GFP fluorescence. TRANS = Brightfield transmitted light, GFP = GFP light source. Error bars represent the ±SD (n = 3). Plaque assay functional titres represent the mean (n = 5).

### 3.4 Replacing the Copper Return Electrode with a Stainless-Steel Electrode indicated that Electrostatic Precipitation was the Sole Cause of Viral Inactivation

In previous runs, copper tape was attached to the positively charged return electrode, functioning as a collector plate for the precipitation of ionised virus particles. However, copper is a naturally virucidal metal and studies have shown direct contact between copper and viral particles resulting in viral inactivation [30]. Therefore, we hypothesised that direct contact between the aerosolised viral particles and the copper tape may have been causing the viral inactivation observed in previous runs. To determine whether EP or the copper tape was causing viral inactivation, stainless-steel sheets were used to replace the copper tape. Stainless-steel is a biologically inert, non-toxic metal [31], and should not inactivate Ad5 particles upon direct contact. Ad5 samples that were not aerosolised, nor exposed to EP, were exposed to the stainless-steel sheets (direct contact for 2 minutes) and analysed for viral activity in the same way as the collected experimental samples.

There was no significant difference between the number of Ad5 viral genomes in the non-exposed Ad5 sample and the Ad5 sample that was exposed to stainless-steel ***(Figure 5.A***). This indicated that stainless-steel did not alter the integrity of the viral DNA. The number of Ad5 genomes was significantly decreased in the Ad5 sample exposed to inactive EP, indicating that aerosolization alone resulted in a reduce in viral DNA collected within the BioSampler. However, the number of viral genomes was further significantly reduced in Ad5 samples following exposure to active EP at 8kV and 10kV (***Figure 5.A***). This indicated that EP successfully captured the aerosolised Ad5 particles. Cells treated with non-exposed Ad5 and the Ad5 sample that was non-exposed to the closed-system but exposed to stainless-steel showed no significant difference in the percentage of virally transduced cells ***(Figure 5.B***). Plaque assay results mirrored this result, showing no visible differences between TREx-293T cells infected with either sample ***(Figure 5. C***). This indicated that direct contact between Ad5 particles and stainless-steel did not affect viral viability. In addition, CHO-CAR cells infected with Ad5 samples exposed to active EP at 8kV and 10kV displayed 11.32-fold and 86.9-fold reductions in the percentage of virally transduced cells, indicating successful inactivation of Ad5 particles by EP ***(Figure 5.B***). Confirming this, TREx-293T cells infected with Ad5 samples that had been exposed to active EP at 8kV and 10kV showed visibly reduced levels of fluorescence, indicating successful inactivation ***(Figure 5.C***).

**Figure 5.**
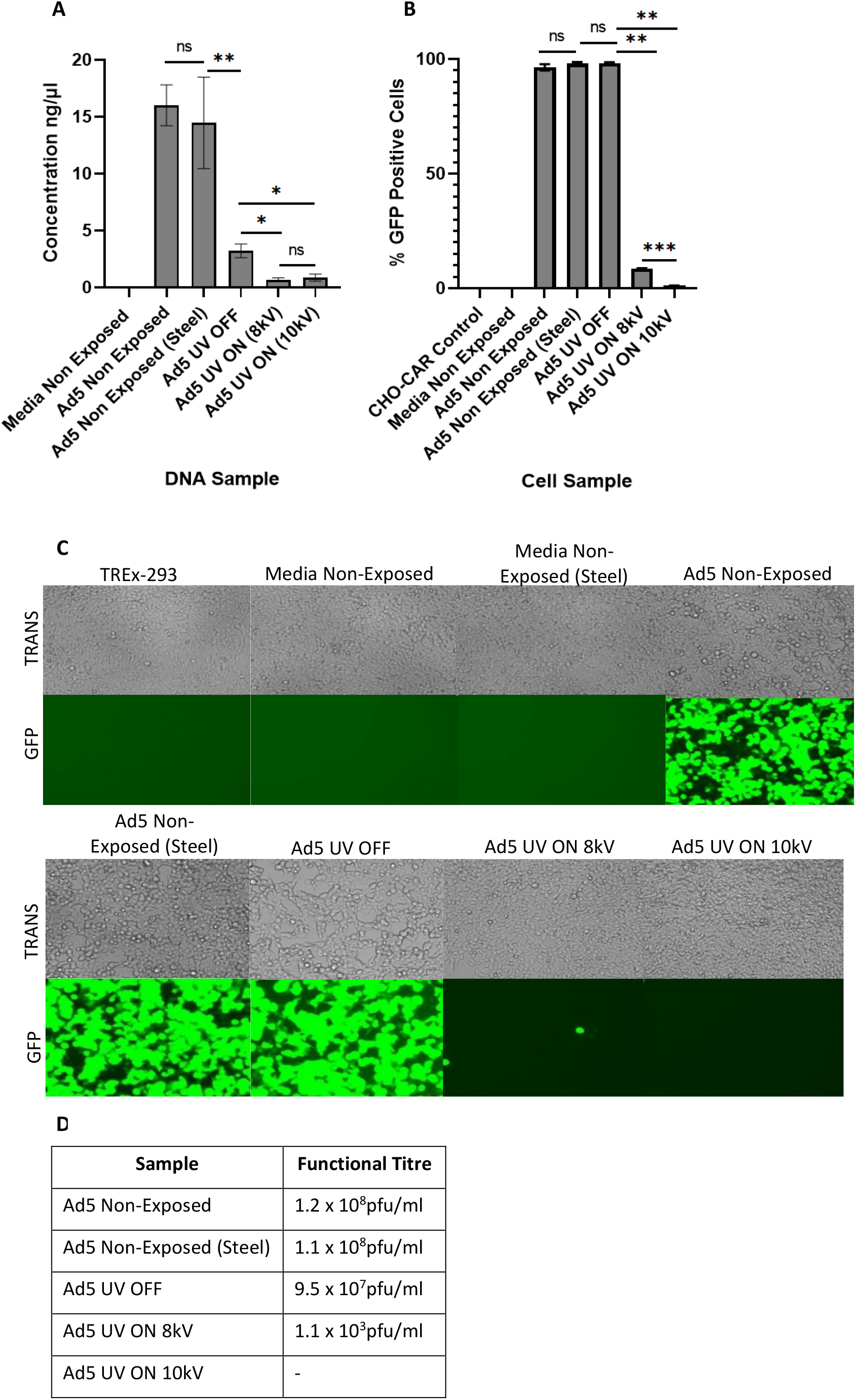
Evidencing EP as the Sole Cause of Viral Inactivation. ‘UV OFF’ signifies sample exposure to inactive Ultravision^TM^ and UV ON’ signifies sample exposure to active Ultravision^TM^. ‘Non-Exposed’ signifies samples that were not aerosolised through the model system, nor exposed to Ultravision^™^. ‘Steel’ signifies samples that were exposed (direct contact) to stainless-steel for 2 minutes. **(A)** Viral capture demonstrated by qPCR. **(B)** Viral inactivation determined by transduction assay. **(C & D)** Viral inactivation displayed by plaque assay in TREx-293 cells. TREx-293 cells treated with samples and analysed for GFP fluorescence. TRANS = Brightfield transmitted light, GFP = GFP light source. Error bars represent the ±SD (n = 3). Plaque assay functional titres represent the mean (n = 5).

### 3.5 EP Successfully Captured and Inactivated Enveloped Viral Particles (SARS-2 PV)

Finally, we sought to evaluate the ability of EP to capture and inactivate enveloped viral particles, such as SARS-CoV-2. As Ad5 is a non-enveloped virus, we used a SARS-CoV-2 Pseudotyped Lentivirus (SARS-2 PV), as its core and genetic material is enclosed by a lipid envelope which expresses the Wuhan Spike protein on its surface, thereby resembling the external structure of wildtype SARS-CoV-2. Neat samples of SARS-2 PV were aerosolised and exposed to EP in the same way as Ad5 in ***Figure 1*.**

SARS-2 PV was significantly captured by EP, as quantified by qPCR ***(Figure 6.A***). A 2.6-fold reduction in the number of viral genomes was observed in the SARS-2 PV sample that had been exposed to active EP, indicating successful virus capture ***(Figure 6.A***). In addition, transduction and plaque assays using the collected samples showed that EP significantly inactivated aerosolised SARS-2 PV particles ***(Figure 6.B, C & D***). CHO-ACE2-TMPRSS2 cells infected with the SARS-2 PV sample that had been exposed to active EP displayed a 27.7-fold reduction in the percentage of viral transduction ***(Figure 6.B***).

**Figure 6.**
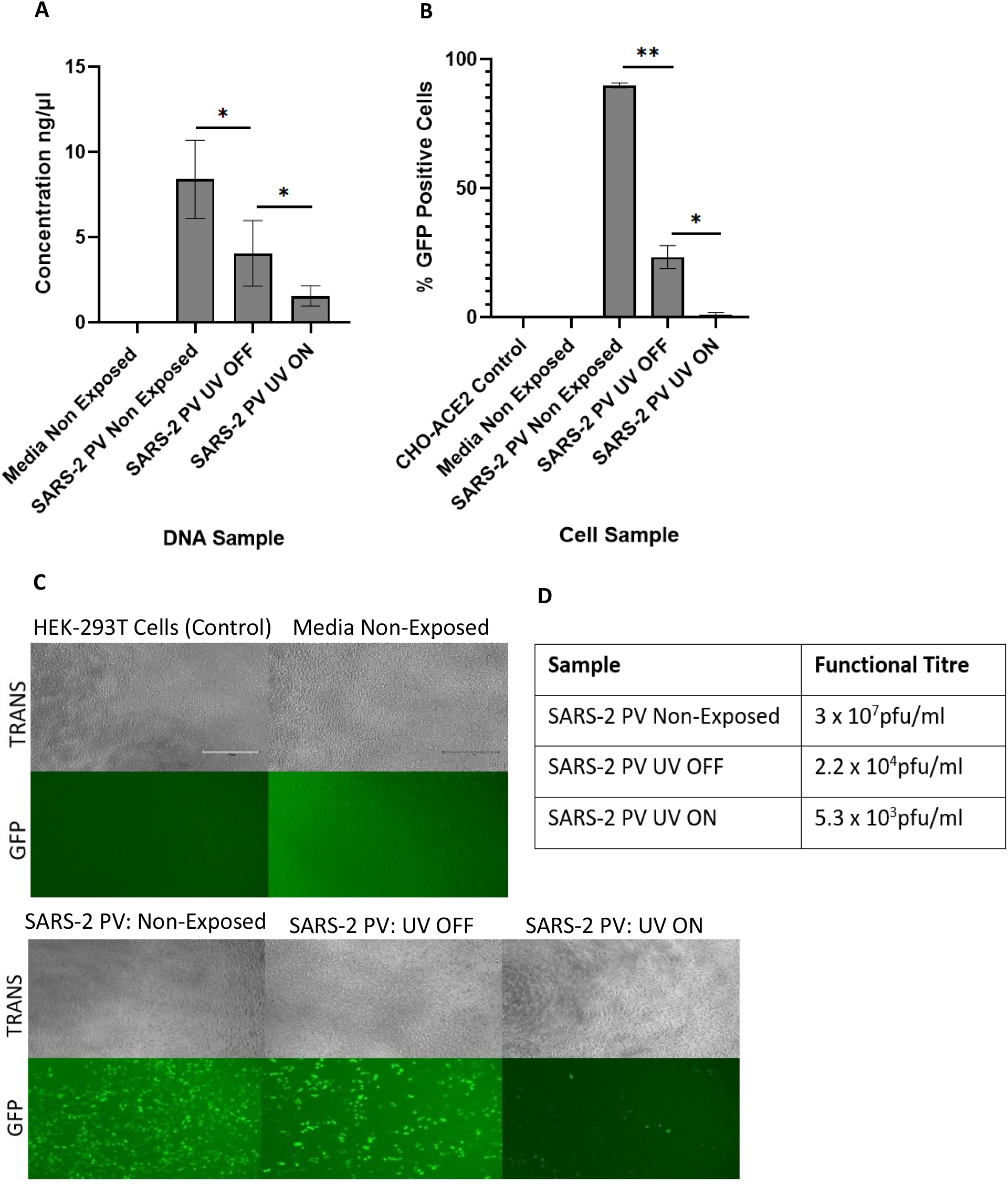
Capture and Inactivation of SARS-2 PV by EP. ‘UV OFF’ signifies sample exposure to inactive Ultravision^™^ and ‘UV ON’ signifies sample exposure to active Ultravision^™^. ‘Non-Exposed’ signifies samples that were not aerosolised through the model system, nor exposed to Ultravision^™^. **(A)** Viral capture determined by qPCR. **(B)** Viral inactivation demonstrated by transduction assay. **(C & D)** Viral inactivation displayed by plaque assay in HEK-293T cells. HEK-293T cells treated with samples and analysed for GFP fluorescence. TRANS = Brightfield transmitted light, GFP = GFP light source. Error bars represent the ±SD (n = 3). Plaque assay functional titres represent the mean (n = 5).

Likewise, HEK-293T cells infected with SARS-2 PV that had been exposed to active EP displayed a visually decreased number of fluorescent cells, compared to the non-exposed sample and the SARS-2 PV sample exposed to inactive EP ***(Figure 6.C***). However, the number of viral genomes, as well as viral viability, was significantly reduced in the SARS-2 PV samples that were aerosolised and exposed to inactive EP (***Figure 6***). This indicated that aerosolised SARS-2 PV was less stable than aerosolised Ad5, and that the sample was more susceptible to inactivation or degradation by aerosolization alone.

## 4. Discussion

Existing methods of purifying indoor air are limited by their inability to capture aerosolised particles smaller than 0.15μm and failure to inactivate live pathogens upon successful capture [12]. These limitations facilitate disease transmission. During periods of viral outbreaks, such as the 2019 SARS-CoV-2 pandemic, bioaerosol-generating medical procedures are at risk of cancellation and delay, due to the likelihood of viral spread [6]. It is therefore crucial that novel non-pharmaceutical interventions (NPI’s) are developed to prevent airborne viral transmission in hospital settings, enabling medical procedures to continue safely and as normal. Established EP systems are currently used to sample and filter indoor air, as well as to clear surgical smoke during key-hole surgeries. Here we have demonstrated additional modalities of the Ultravision^™^ EP device, in its ability to efficiently capture and inactivate aerosolised viral particles.

Significant capture and inactivation of aerosolised Ad5 and SARS-2 PV particles by EP was observed in our standardised closed-system model. Viral capture was displayed by a reduction in the number of viral genomes collected within the BioSampler, following sample exposure to active EP, compared to recovered samples exposed to inactive EP. Similarly, viral inactivation was shown by a reduction in biological activity of viral particles, as gauged by the percentage of transduced cells that were treated with recovered samples post exposure to active EP, compared to samples exposed to inactive EP. Interestingly, it appeared that viral inactivation by EP was more successful than viral capture.

Viral inactivation by EP was highly efficient, at approximately 90-95% efficiency when using Ultravision^™^ at 8kV, and at >99% efficiency when using Ultravision^™^ at 10kV or when using 2 Ultravision^™^ Ionwands (both at 8kV). Arguably, viral inactivation is more important than viral capture, as this can prevent the spread of disease. Previous studies evaluating the ability of EP to inactivate viruses suggest that the corona discharge, produced by the discharge electrode, generates reactive species (O_3_ and various radicals, such as O·, N·, OH·, and HO_2_ ·) capable of degrading and inactivating viral particles [32–34]. Although this mechanism has not been explicitly investigated here, our results indicate that this could be the cause of viral inactivation. In agreement, degradation of viral particles would result in the release of viral DNA/RNA, explaining the collection of viral genomes in the BioSampler following sample exposure to active Ultravision^™^. As isolated viral DNA is biochemically inert and requires an intact capsid to bind and enter target cells, the degradation of aerosolised viral particles seems a practical way of inactivating viruses and reducing their transmission [35, 36].

We have demonstrated that EP can efficiently capture and inactivate both non-enveloped (Ad5) and enveloped (SARS-2 PV) viral particles. However, aerosolization alone significantly reduced SARS-2 PV viability and the integrity of its capsid, causing the release of its viral genome. This was not surprising as SARS-2 PV is not a respiratory virus and is therefore not transmissible via airborne routes. However, other non-respiratory viruses, such as wildtype HIV and HPV, have been identified in surgical bioaerosols with the ability to infect healthcare staff. Therefore, it is important that EP can capture and inactivate a variety of viral particles [3, 4]. Future studies will focus on evaluating the ability of EP to capture and inactivate respiratory enveloped viruses, as well as non-respiratory non-enveloped viruses. In addition, other physical parameters govern viral spread and stability, including temperature, humidity, droplet size and air-space volume [37]. Evaluating changes to viral capture and inactivation, following the alteration of such parameters, as well as parameters effecting the efficiency of EP, such as voltage, flow rate, geometric design of the EP system and size and concentration of the ionised particles [38], will be important to optimise in future studies, prior to implementing EP in hospitals as a method of reducing viral spread.

Regarding Ultravision^TM^ as a multi-model device, it may have a role to play beyond key-hole surgery. Due to recent advances in Ultravision^™^ technology, it is likely that Ultravision^™^ will be employed during open surgeries in the near future to clear surgical smoke. It is therefore possible that Ultravision^™^ could be manipulated to capture and inactivate viral particles in ‘open’ systems. However, this would of course require adaptations to the device itself to enable sufficient exposure of the corona discharge to bioaerosols covering a much larger surface area upon release from the patient. As well as this, Ultravision^™^ could be implemented when delivering aerosolised medications to patients. For example, pressurised intraperitoneal aerosol chemotherapy (PIPAC) has recently been developed as a method of treating unresectable metastatic peritoneal tumours [39, 40]. Using EP in this way would ensure efficient delivery of drugs or oncolytic viruses by PIPAC to the tumour site, whilst simultaneously enhancing the safety of its administration, preventing escape of the drug or therapeutic ATMPs into the surrounding environment.

In summary, our findings indicate that EP could be used during surgery to capture and inactivate viral particles released in bioaerosols, as well as potentially during other medical procedures, to enhance efficacy and safety. Employing EP as a NPI to reduce viral spread in hospitals may resolve issues experienced with existing air-purification systems, which in turn could reduce pressures on the NHS by preventing indirect morbidities and mortalities. For example, recent outbreaks of the Highly Pathogenic Avian Influenza A(H5N1) in wild birds and poultry has the capacity to spread to human hosts, which if unprevented, could result in the next human global pandemic [41]. Using data obtained from this study, we predict that it is possible to use EP to minimise viral spread thus preventing future viral pandemics.

## Supporting information

Supplementary figures, Preston HE et al.

## Acknowledgements

We thank Dominic Griffiths (Alesi Surgical Ltd) for helpful discussions on this project. We also thank Michael Shinkwin and Neil Warren for helping to build an early prototype model for this study.

