## Supplementary figures, Preston HE et al. for "Efficient Viral Capture and Inactivation from Bioaerosols Using Electrostatic Precipitation"

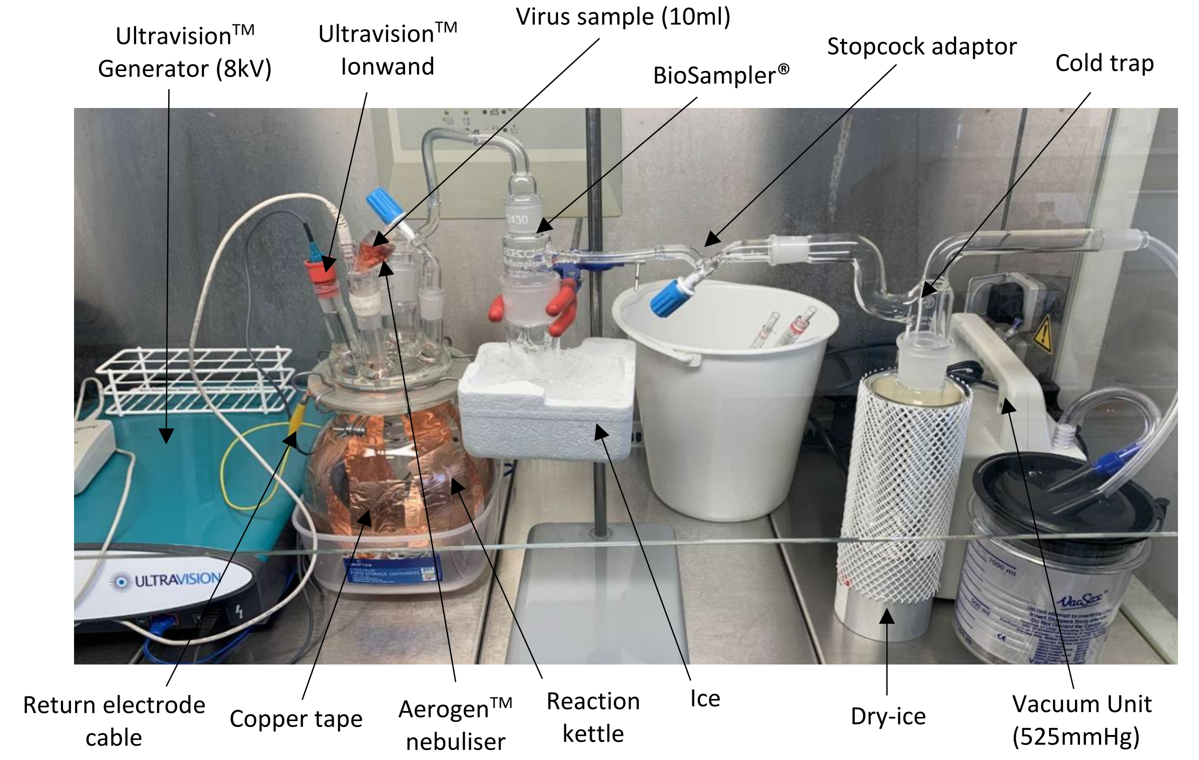


***Supplementary Figure 1.*** *Experimental Setup of the Refined Closed-System Model. All samples were aerosolised into the air-tight reaction kettle, exposed to Ultravision™ (active/inactive) and suctioned into the BioSampler for recovery and collection. Collected samples were stored at -80°C immediately after each experimental run, prior to experimental analysis.*

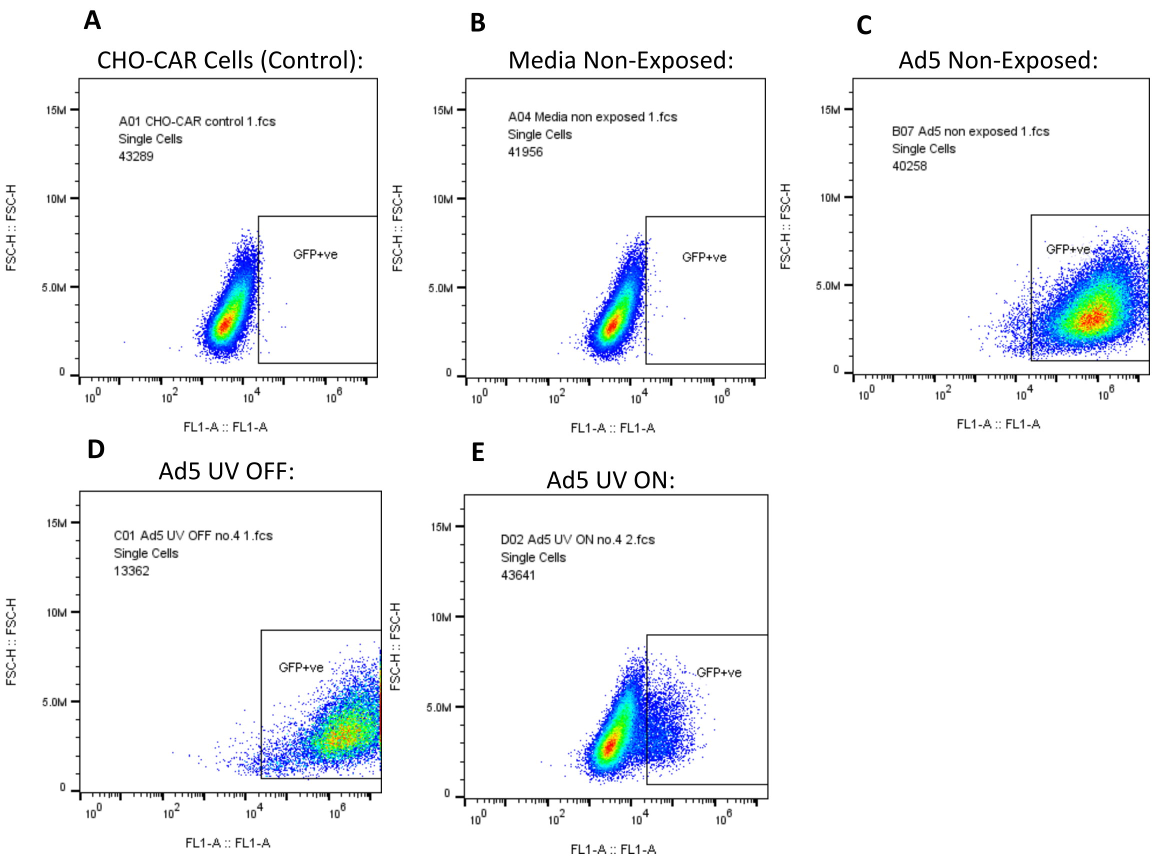


***Supplementary Figure 2.*** *Transduction assay raw data (**Figure 2.B**). Cell populations infected with experimental samples analysed by Flow Cytometry and gated with the FL1-A channel, to detect transduced (GFP positive) cells.*

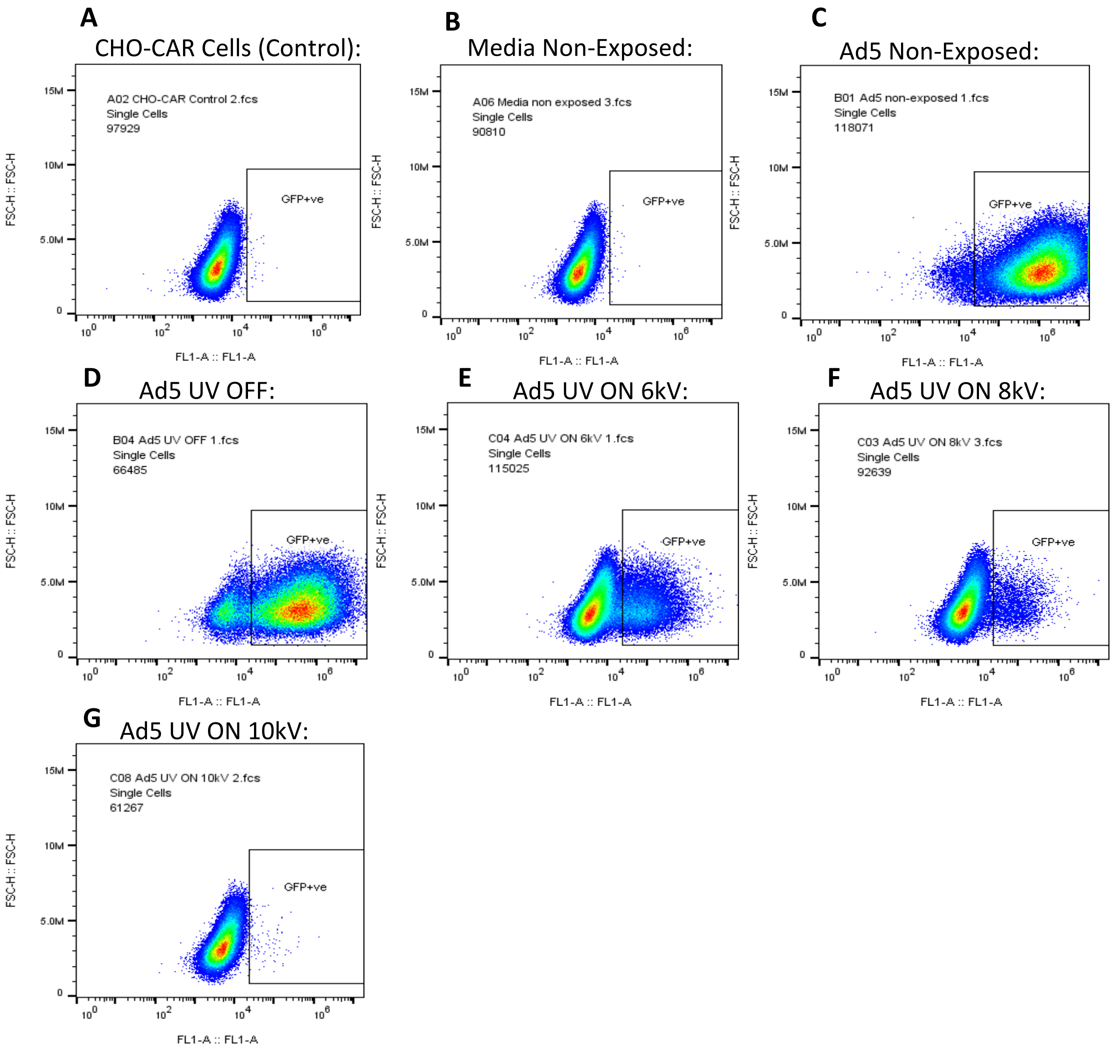


***Supplementary Figure 3.*** *Transduction assay raw data (**Figure 3.B**). Cell populations infected with experimental samples analysed by Flow Cytometry and gated with the FL1-A channel, to detect transduced (GFP positive) cells.*

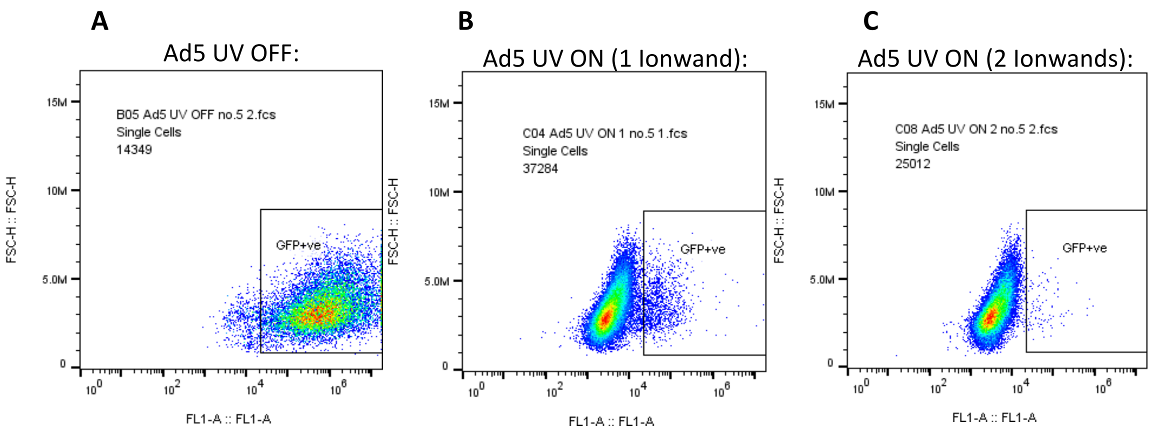


***Supplementary Figure 4.*** *Transduction assay raw data (**Figure 4.B**). Cell populations infected with experimental samples analysed by Flow Cytometry and gated with the FL1-A channel, to detect transduced (GFP positive) cells.*

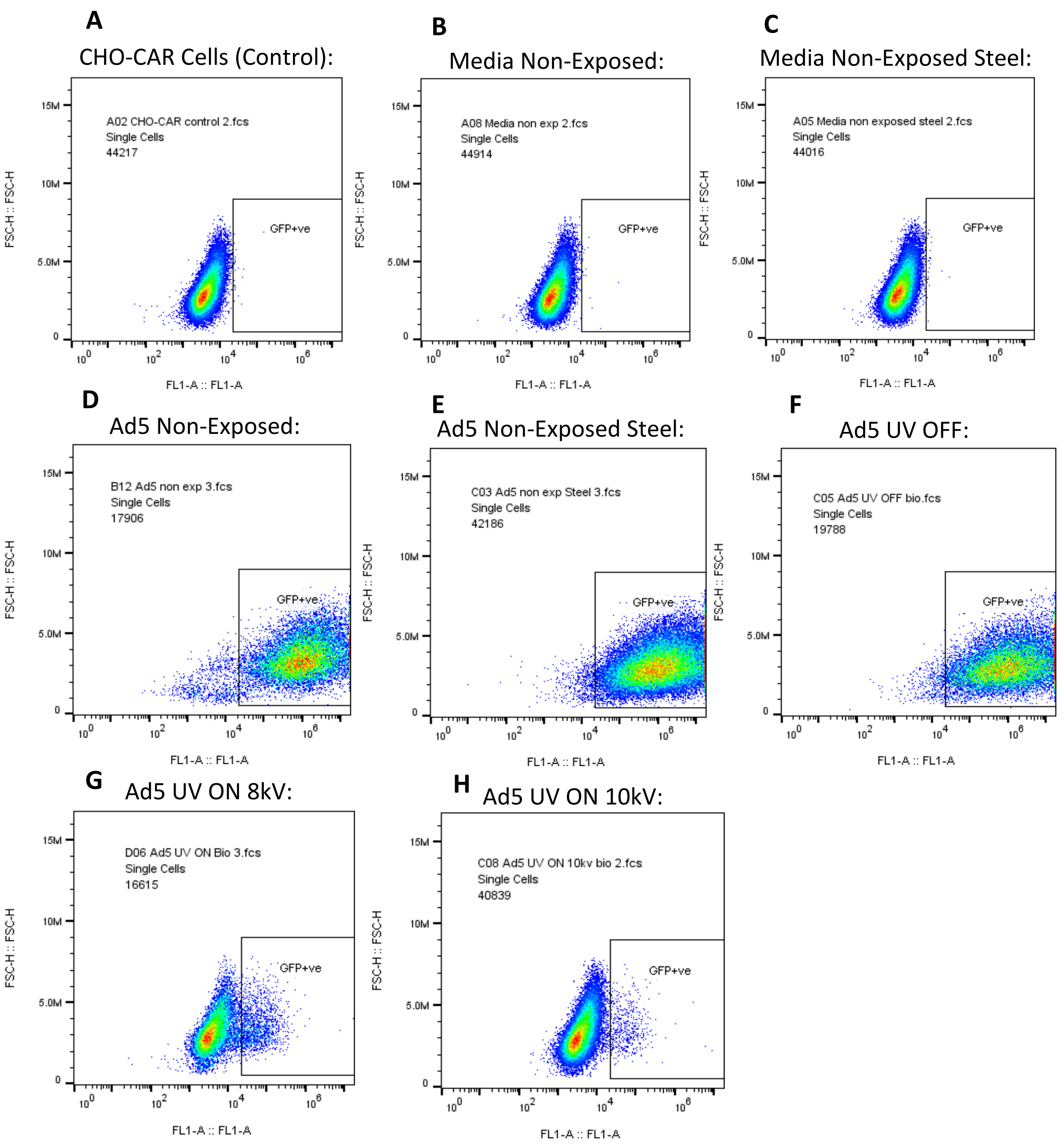


***Supplementary Figure 5.*** *Transduction assay raw data (**Figure 5.B**). Cell populations infected with experimental samples analysed by Flow Cytometry and gated with the FL1-A channel, to detect transduced (GFP positive) cells.*

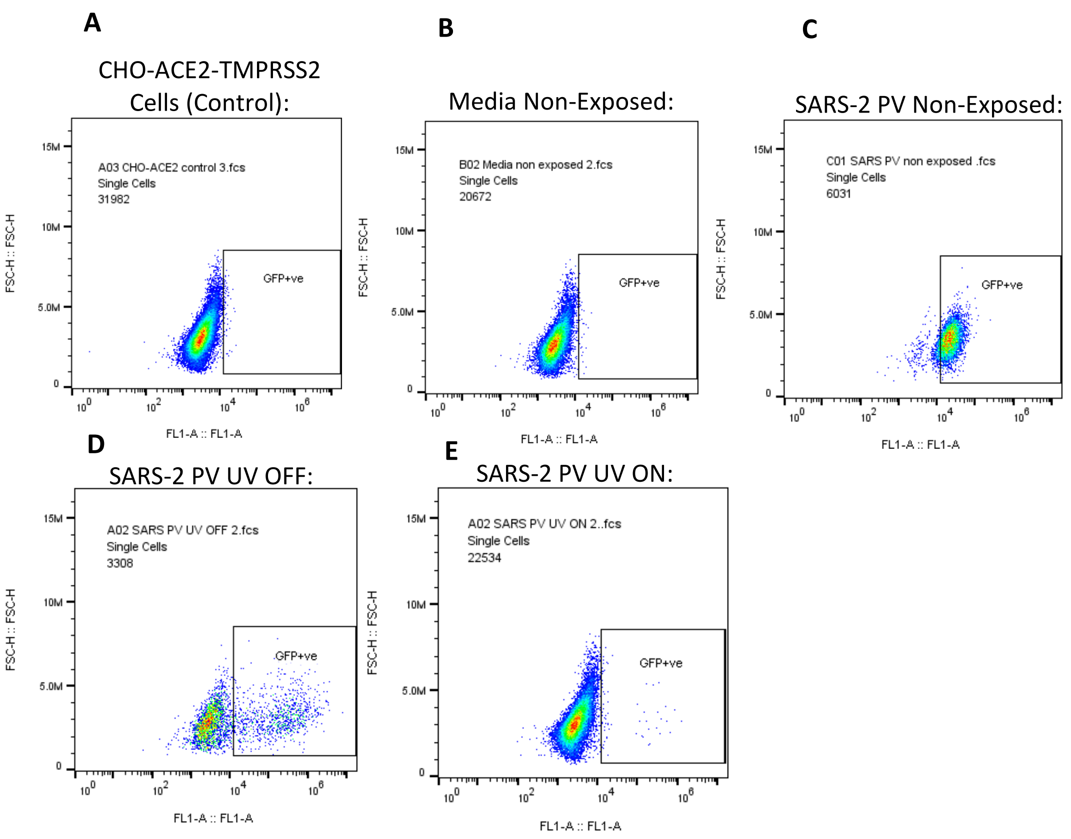


***Supplementary Figure 6.*** *Transduction assay raw data (**Figure 6.B**). Cell populations infected with experimental samples analysed by Flow Cytometry and gated with the FL1-A channel, to detect transduced (GFP positive) cells.*
